# Comparative pangenomic insights into the distinct evolution of virulence factors among grapevine trunk pathogens

**DOI:** 10.1101/2023.09.03.555958

**Authors:** Jadran F. Garcia, Abraham Morales-Cruz, Noé Cochetel, Andrea Minio, Rosa Figueroa-Balderas, Philippe E. Rolshausen, Kendra Baumgartner, Dario Cantu

**Affiliations:** Department of Viticulture and Enology, University of California, Davis, Davis, CA, United States; U.S. Department of Energy, Joint Genome Institute, Lawrence Berkeley National Lab, Berkeley, CA, USA; Department of Botany and Plant Sciences, University of California, Riverside, Riverside, CA, United States; Crops Pathology and Genetics Research Unit, United States Department of Agriculture – Agricultural Research Service, Davis, CA, United States; Genome Center, University of California, Davis, Davis, CA, United States

## Abstract

The permanent organs of grapevines (*V. vinifera* L.), like other woody perennials, are colonized by various unrelated pathogenic ascomycete fungi secreting cell wall-degrading enzymes and phytotoxic secondary metabolites that contribute to host damage and disease symptoms. Trunk pathogens differ in the symptoms they induce and the extent and speed of damage. Isolates of the same species often display a wide virulence range, even within the same vineyard. This study focuses on *Eutypa lata*, *Neofusicoccum parvum*, and *Phaeoacremonium minimum*, causal agents of Eutypa dieback, Botryosphaeria dieback, and Esca, respectively. We sequenced fifty isolates from viticulture regions worldwide and built nucleotide-level, reference-free pangenomes for each species. Through examining genomic diversity and pangenome structure, we analyzed intraspecific conservation and variability of putative virulence factors, focusing on functions under positive selection, and recent gene-family dynamics of contraction and expansion. Our findings reveal contrasting distributions of putative virulence factors in the core, dispensable, and private genomes of each pangenome. For example, CAZymes were prevalent in the core genomes of each pangenome, whereas biosynthetic gene clusters were prevalent in the dispensable genomes of *E. lata* and *P. minimum*. The dispensable fractions were also enriched in Gypsy transposable elements and virulence factors under positive selection (polyketide synthases genes in *E. lata* and *P. minimum* glycosyltransferases in *N. parvum*). Our findings underscore the complexity of the genomic architecture in each species and provide insights into their adaptive strategies, enhancing our understanding of the underlying mechanisms of virulence.

## Introduction

Eutypa dieback, Botryosphaeria dieback, and Esca are widespread and damaging trunk diseases, which decrease grapevine (*Vitis vinifera* L.) productivity and longevity (1, 2). *Eutypa lata, Neofusicoccum parvum,* and *Phaeoacremonium minimum* are among the most widely distributed and virulent species associated with Eutypa dieback, Botryosphaeria dieback, and Esca, respectively. These fungi represent different orders within the Fungal Phylum Ascomycota: *E. lata* (Class Sordariomycetes, Order Xylariales, Family Diatrypaceae), *N. parvum* (Class Dothideomycetes, Order Botryosphaeriales, Family Botryosphaeriaceae), and *P. minimum* (Class Sordariomycetes, Order Togniniales, Family Togniniaceae). They infect grapevines through wounds, colonizing permanent woody structures, such as trunks, cordons, and spurs. Fungal colonization disrupts the distribution of water and nutrients within the vine, resulting in the death of shoots and woody tissues. These diseases are characterized by external symptoms on the leaves and fruit, and internal symptoms in the wood (Figure 1).

**Figure 1.**
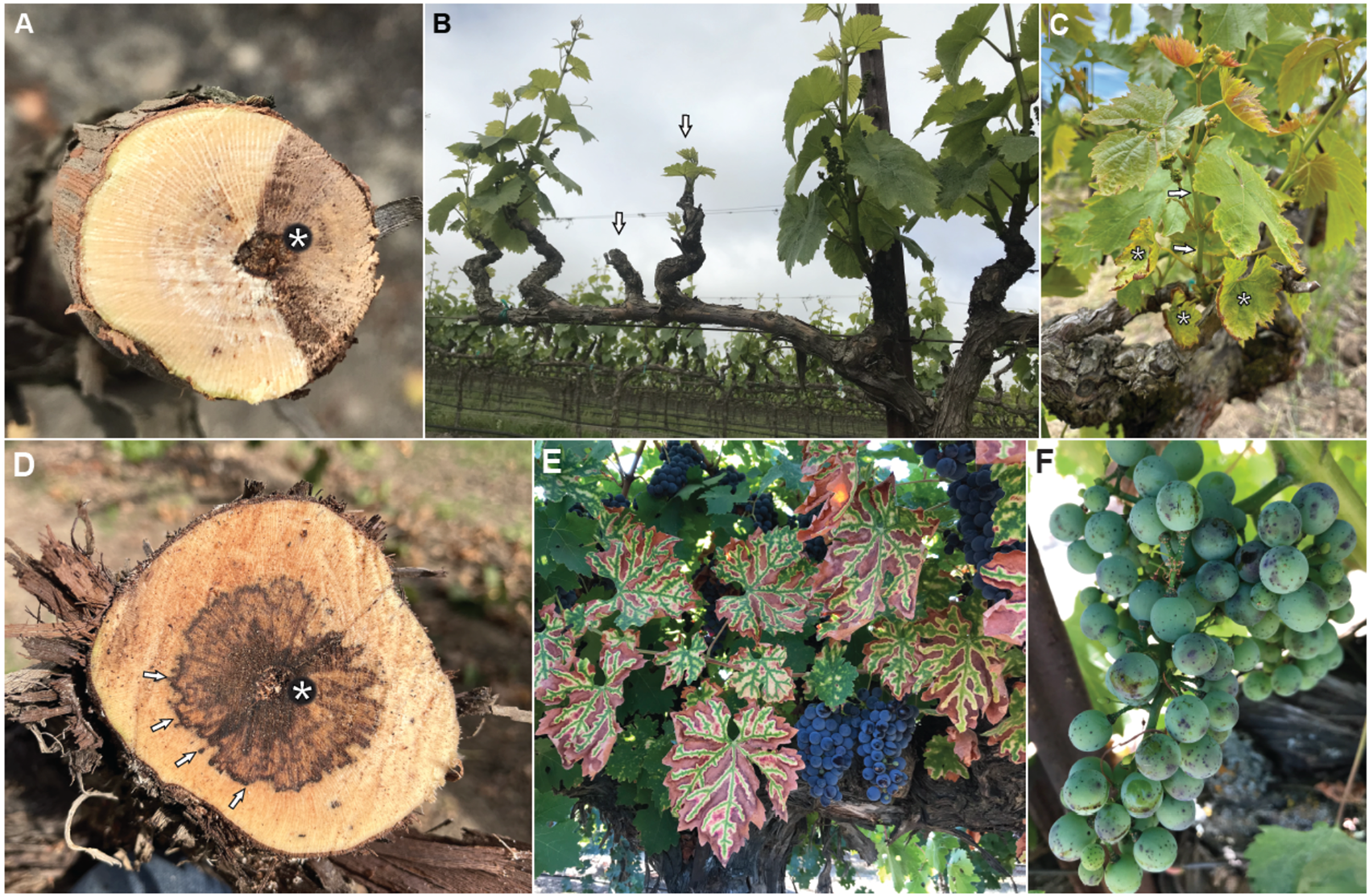
Common symptoms of the grapevine trunk pathogen species in this study. This figure shows different pictures of the common wood and foliar symptoms of the diseases in this study. **A**, Cross-section showing wood canker: a necrotic, wedge-shaped discoloration characteristic of Botryosphaeria dieback and Eutypa dieback. **B**, Cordon of a vine showing stunted shoots, and dead spur, characteristic of Botryosphaeria dieback (arrows). **C**, Shoot of a vine with stunted growth (short internodes, arrows). The leaves are small, cupped, and chlorotic with necrotic margins (**\***), characteristic of Eutypa dieback . **D**, Cross-section showing central vascular streaking (**\***), surrounded by black spots and vascular streaking (arrows), characteristic of Esca. **E**, Common interveinal discoloration and scorching characteristic of Esca. **F**, Common Esca fruit symptoms showing dark spots on berries (also known as measles).

The destructive colonization of woody tissues hinges on several virulence factors, including enzymes that degrade cell walls, toxins, and a variety of cellular transporters (3–7). *E. lata*, for example, breaks down cellulose and other cell-wall glycans, also secreting oxidases that may degrade wall-bound lignin in the wood (8). Various secondary metabolites with phytotoxic properties, such as eutypine, eutypinol, scytalone, isosclerone, mellein, tyrosol, and others, have been detected in the secretomes of pathogens causing Eutypa, Esca, and Botryosphaeria dieback (9–12). Even if the precise mode of action of these molecules is not fully understood, it is known that fungal toxins can harm plant cells by interfering with enzymatic reactions, cellular transport, and causing damage to cell membranes (6, 13). Further damage is associated with the host response to infection, namely the formation of xylem vessel occlusions (14, 15).

In addition to their importance for the grape industry, *E. lata*, *N. parvum*, and *P. minimum* serve as valuable models for studying the mechanisms of pathogenicity and virulence-factor evolution in phylogenetically unrelated species that colonize the same cell and tissues. One of the reasons for their significance lies in the fact that although all three species colonize the same woody tissue, they differ in terms of virulence and the specific disease symptoms they induce. This diversity implies a distinct deployment of virulence-factor repertoires by each species. Early comparisons of genome assemblies indicated varied gene-family expansions and contractions associated with virulence factors in *E. lata*, *N. parvum*, and *P. minimum* (3). Notably, species eliciting similar symptoms were reported to share common gene-family expansions, indicating possible convergent evolution, irrespective of phylogenetic relationships. For example, Esca pathogens *P. minimum* and *Phaeomoniella chlamydospora*, despite the fact that they represent different Classes of Fungi, showed similar sugar-transport expansion. Similarly, *E. lata*, *N. parvum*, and Phomopsis-dieback pathogen *Diaporthe ampelina*, all of which cause similar wood cankers, had comparable expansions in specific families of CAZymes, polyketide synthases, and P450s.

Furthermore, within each of the three species, there is a significant degree of variability in virulence among isolates. For example, a study on *E. lata* isolates inoculated on grapevines showed notable differences in both the severity of wood lesions (16, 17) and the incidence of foliar symptoms (17). Similar observations were made with *N. parvum*, where significant intraspecific virulence variations were observed in grapevine green shoots and potted plants from different regions of New Zealand, Australia, South Africa, and California (18). Interestingly, for both species, even isolates from the same vineyard showed differences in virulence (16, 17). *P. minimum* also exhibited significant virulence variability, as evident from variations in lesion length and dry shoot weight among grapevines inoculated with different isolates (19).

The broad variability in virulence observed among isolates is likely due to differences in virulence-factor content, properties, and expression during infection (5, 20). For instance, in the case of *P. minimum*, genomes of different isolates exhibit approximately 1 Mbp of structural differences, resulting in the presence/absence polymorphisms of multiple adjacent genes, particularly associated with biosynthetic clusters related to secondary metabolism (5). Indels also impact complete genes responsible for the biosynthesis of polyketides, a diverse group of biologically active compounds found in fungi, such as mycotoxins, antifungal, and antibiotic products. Comparable findings were reported in *E. lata*, where structural variants and chromosomal rearrangements were linked to alterations in secondary-metabolite production (20).

Consequently, a single genome reference fails to represent the complete virulence factor repertoire of a species, as demonstrated in the case of *P. minimum* (5). Using only one reference genome has been shown to limit the effectiveness of identifying intraspecific genetic variations, particularly novel or highly divergent sequences present within a species (21, 22). Pangenomes have been developed to provide a comprehensive view of the genetic landscape that may otherwise be overlooked (23–25). Pangenomes identify genomic segments shared by all genomes in a species, as well as those shared by only some isolates or exclusively present in individual isolates. (22, 23). In fungal plant pathogens, conserved (‘core’) genomic regions were described to mainly consist of ‘house-keeping genes’, with a few virulence factors believed to be essential for pathogenesis, whereas the variable (‘dispensable’) and unique (‘private’) regions are often enriched in virulence factors (23, 26, 27). These virulence factor-rich regions, present in dispensable and private genomic regions, serve as strong evidence in support of the differences in virulence among isolates. Therefore, using pangenomes in species representation provides a more comprehensive understanding of the evolutionary patterns of these factors within a species. Examining multiple pangenomes can shed light on species-specific evolutionary trends (28).

Following this rationale, in this study we produced nucleotide-level, reference-free pangenomes for *E. lata*, *N. parvum*, and *P. minimum* utilizing fifty *de novo* assembled genomes from isolates collected across a wide range of geographical locations. We examined genomic diversity and pangenome structure to analyze the intraspecific conservation and variability of virulence factors, with an emphasis on functions under positive selection and the dynamics of recent gene-family changes. By comparing the three pangenomes, we highlight interspecific differences in virulence function evolution potentially reflecting species-specific virulence strategies and differing evolutionary pressures.

## Results

### Assembly of fifty fungal genomes

To construct comprehensive pangenomes for the three species, we *de novo* assembled the genomes from multiple isolates—six for *P. minimum*, 16 for *N. parvum*, and 28 for *E. lata*. These isolates were sampled from grape-producing regions across North and South America, Europe, Africa, and Oceania, to ensure a broad representation of genomic diversity (Figure 2, Supplementary table 1). The countries sampled included USA, Chile, France, Spain, Portugal, Greece, South Africa, and Australia. For some countries, we were able to include isolates from distinct viticultural regions, enhancing the geographical coverage and genomic representation.

The fifty genomes were short-read sequenced at an average coverage of 93 ± 6 X. The total size of the *de novo* assembled genomes was 46.5 ± 0.3 Mbp for *P. minimum* (41.0 ± 2.1 Mbp, estimated genome size based on kmer distribution (29, 30)), 43.5 ± 0.2 Mbp for *N. parvum* (44.4 ± 0.2 Mbp, estimated size), and 54.9 ± 0.1 Mbp for *E. lata* (57.3 ± 0.4 Mbp, estimated size). Genome completeness was 97.98 ± 0.80% for *P. minimum*, 98.58 ± 0.05% for *N. parvum*, and 97.50 ± 0.03% for *E. lata*, as evaluated by BUSCO v.5.4.2 (Figure 2). Because genome completeness is essential for pangenome construction (31, 32), we further verified the assembly completeness by comparing the genome assemblies based on short-reads with chromosome-scale assemblies produced through long-read sequencing (Supplementary table 1). We have previously reported the chromosome-scale assemblies of the genomes of *P. minimum* isolate Pm1119 (5) and *N. parvum* isolate UCD646So (6). To generate a genome reference for *E. lata*, we resequenced the DNA of the isolate EN209 using Single Molecule Real-Time (SMRT) technology. Reads were assembled into 213 polished contigs, with a 128 X mean coverage and a total size of 55.9 Mbp. The 10 longest contigs comprised 54.5 Mbp, approximately 98% of the total assembly size and approximately 200 kbp more than the estimated genome size (54.3 Mbp). The remaining unplaced 1 Mbp did not contain any protein-coding genes and were highly repetitive (21 %). The eleventh-longest contig of the assembly corresponded to the complete mitochondrial genome, with a size of 179,645 bp and a coverage of 1,652 X. These findings suggest that the 10 largest contigs represent complete, telomere-to-telomere chromosomes: (i) all 10 contigs contain telomeric repeats at both extremes; (ii) they contain 99.65% of the complete BUSCO genes; and (iii) they have comparable length (3.4 – 7.4 Mbp) to complete fungal chromosomes (33–35). Using these chromosome-scale assemblies, we determined that an average of 89 ± 5% of the genes in *P. minimum* are in collinear blocks of at least five genes, compared to 94 ± 0.6 % in *E. lata* and *N. parvum*. Likewise, an average of 96 ± 1% of the genes in the *P. minimum* reference genome were covered by the short reads assemblies, compared to 97 ± 0.1%, and 98 ± 0.3% in *E. lata* and *N. parvum*, thereby corresponding to the results of BUSCO.

### Virulence factor-focused annotation of the fifty fungal genomes

Functional annotations of the predicted protein-coding genes were conducted with a focus on functions associated with the mechanisms of virulence, utilizing various specialized databases. Specifically, we employed:

(i) AntiSMASH (36) to identify Biosynthetic Gene Clusters (BGCs), which are responsible for the biosynthesis of secondary metabolites. Some of these metabolites can exert phytotoxic effects (37, 38).
(ii) The Fungal P450 database (39) to annotate Cytochrome P450 enzymes. While some are part of BGCs, they also perform various functions such as xenobiotic compound detoxification and primary metabolite production, contributing to pathogenicity (40–42).
(iii) Carbohydrate active enzymes (CAZymes) database to annotate secreted cell wall modifying enzymes.
(iv) Transporter Classification Database (TCDB; (43)) to annotate cellular transporters, which can contribute to host colonization by secretion of numerous molecules, including toxins and enzymes (44).

We identified 14,452 ± 166, 13,030 ± 24, and 15,164 ± 15 predicted protein-coding genes in *P. minimum*, *N. parvum*, and *E. lata*, respectively (Supplementary table 2). The repetitive content was low, as expected, with 1.79 ± 0.10% for *P. minimum*, 5.17 ± 0.07% for *N. parvum*, and 4.92 ± 0.08% for *E. lata* (Supplementary table 2). On average, across all fifty genomes, we found 70 ± 2 BGCs, 960 ± 8 Cytochrome P450 enzymes, and 614 ± 8 CAZymes, half of which were classified as secreted by signalP v5.0 (45). Additionally, an average of 2,976 ± 49 genes were identified as potential cellular transporters. Specific differences were noted among the species confirming what reported in (3), such as *E. lata* having 82 ± 1 BGCs, more than *P. minimum* (42 ± 1) and *N. parvum* (58 ±1). The proportion of P450 enzymes and CAZymes varied, with *E. lata* and *N. parvum* having higher number of P450 enzymes, and *N. parvum* having the highest number of secreted CAZymes. Transporter genes were more abundant in *P. minimum* and *N. parvum* compared *to E. lata* (Supplementary table 2).

**Figure 2.**
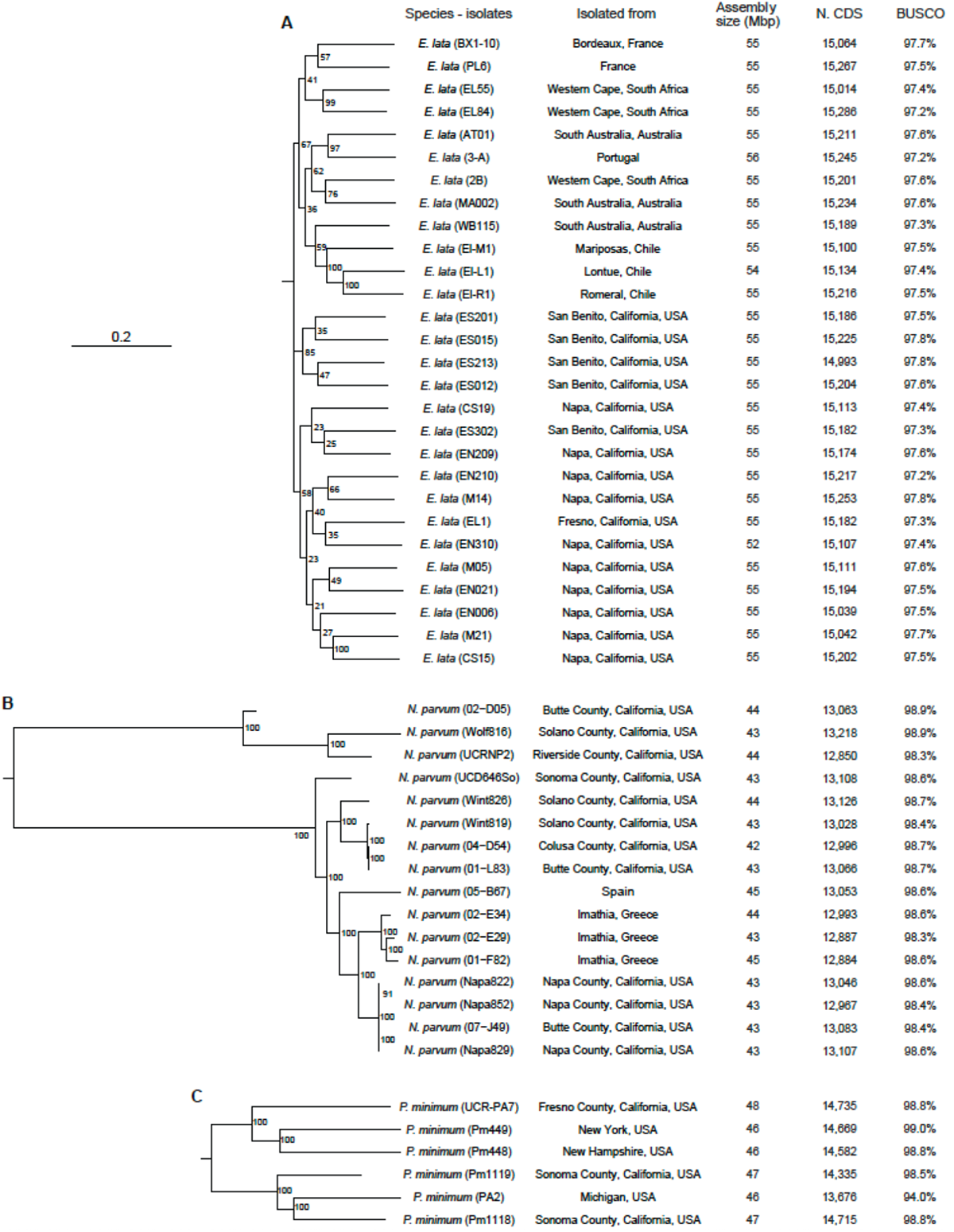
Phylogenetic relationship, origin and summary statistics of the isolates analyzed. The figure includes Maximum likelihood phylogenetic trees with the regions of origin and genome assembly statistics of the isolates form **A**, *Eutypa lata*. **B**, *Neofusicoccum parvum*, and **C**, *Phaeoacremonium minimum*. When genomes from different countries were available, some regional structuring patterns were observed. In *N. parvum*, the isolates from Greece were clustered together. In *E. lata*, all the isolates from the USA formed a single distinct cluster, and isolates from Chile were grouped with one isolate from Australia. Similarly, isolates from South Africa either clustered with Australian or French isolates.

### Construction and analysis of pangenome graphs reveal distinct genomic structures and complexities among the three species

We built for each species a nucleotide-level reference-free pangenome using the tools that are part of the PanGenome Graph Builder (PGGB; (46)) pipeline. The pangenome graph for *P. minimum* consisted of over 4 million nodes, totaling 59.8 Mbp in length (Figure 3A), representing approximately 1.3 times the genome size of the species. Similarly, the pangenome graph for *N. parvum* contained a comparable number of nodes, with a final size of 56.4 Mbp, also about 1.3 times the genome size of the species (Figure 3A). In contrast, the pangenome graph for *E. lata* exhibited a more complex structure, comprising more than 13 million nodes and extending to 89.7 Mbp in length (Figure 3A), roughly 1.6 times the size of an individual genome.

In the pangenome graphs, nodes were categorized as “core” if present in all genomes, “dispensable” if found in more than one (but not all) genomes, and “private” if exclusive to one genome within the pangenome. The largest fraction of nodes in the three pangenome graphs were classified as dispensable: 46.6% in *P. minimum*, 55.6% in *N. parvum*, and 57.2% in *E. lata*. Despite this, the core nodes constituted the largest portion of the pangenome graph length, accounting for 65.9%, 67.0%, and 48.4% in *P. minimum*, *N. parvum*, and *E. lata*, respectively.

Regarding the total pangenome graph length, *Eutypa lata* exhibited the greatest proportion of variable regions in its pangenome, with 51.6% of the total length of the graph being either private or dispensable. This contrasts with *P. minimum* and *N. parvum*, where only 34.2% and 33.0% of the total graph length were variable (Figure 3A).

Repetitive regions constituted a larger proportion of the private sequence length in all three species (Figure 3B). In *P. minimum*, 8.1 ± 0.1% of the private genome was composed of repeats, in contrast to 4.0 ± 0.3% and 1.3 ± 0.01% in the dispensable and core regions, respectively. *E. lata* followed a similar pattern, with 2.7 ± 0.01% of the core genome covered by repeats, compared to a significantly higher 28.2 ± 1.5% in the private genome (Wilcox test, *P* = 6.65e-17). The most pronounced difference was observed in *N. parvum*, where a substantial 40.0 ± 5.0% of the private genome consisted of repetitive regions. Additionally, among the transposable elements detected, the LTR/Gypsy were the most abundant with 6.1 ± 0.7% in *E. lata*, 11.5 ± 2.6% in *P. minimum* and 12.4 ± 1.2% in *N. parvum*. A significant over-representation of Gypsy elements (*P* < 0.05, Wilcox test) was found in the private genomes of the three species (Supplementary figure 1 and 2), suggesting that these elements in particular play an important role in genome evolution in these species.

**Figure 3.**
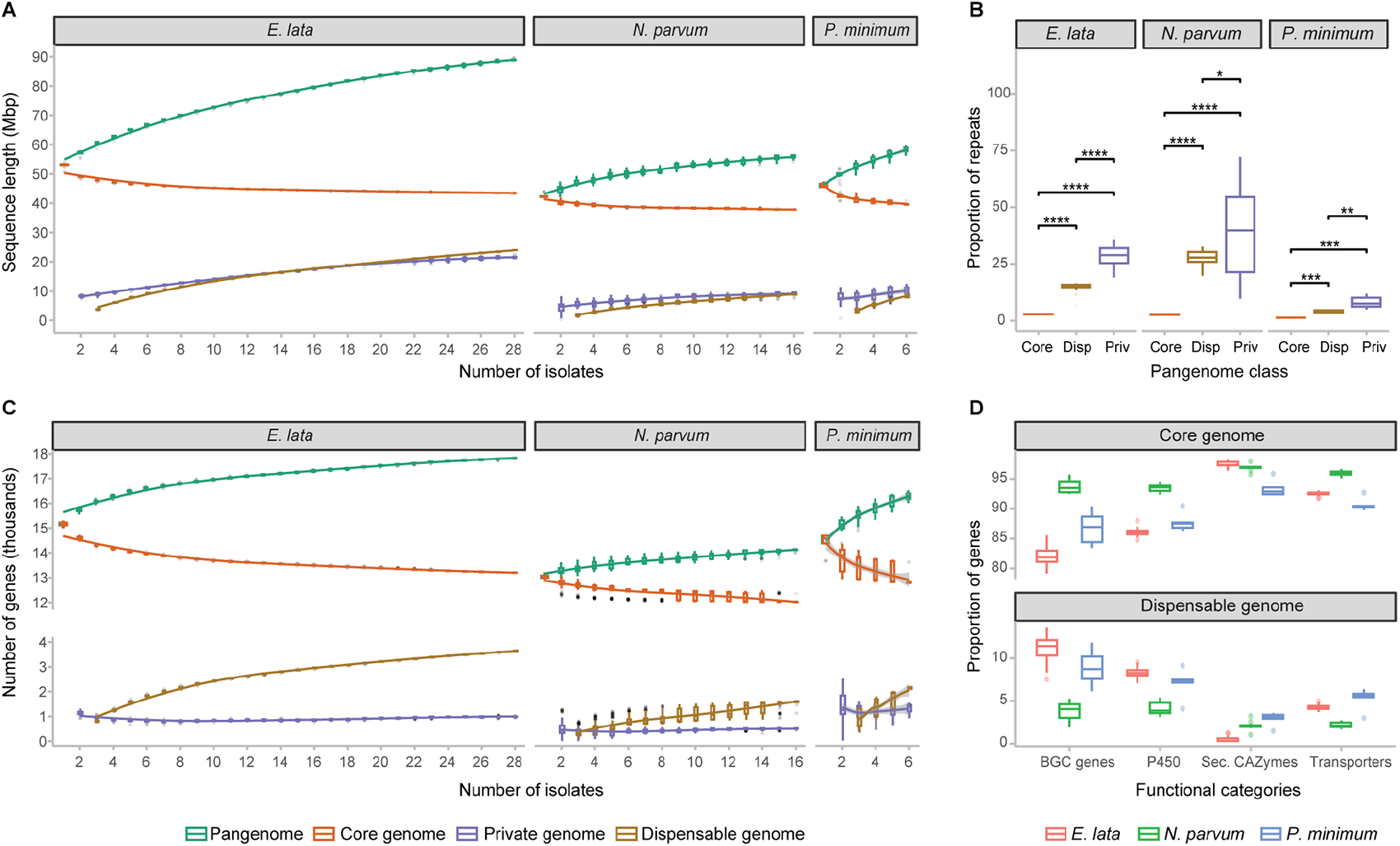
Properties of the seq-based and gene-based pangenome. **A**, Sequence-based pangenome modeling. For each combination of genomes, the total length of the nodes is represented per class. The line represents smoothed conditional means with a 0.95 confidence interval. **B**, Proportion of the different node classes annotated as repeats. Disp means Dispensable and Priv means Private. Significant differences between groups were determined after a Kruskal-Wallis test (*P* < 0.05). **C**, Gene-based pangenome modeling. For each combination of genomes, the number of genes is represented per class. The lines represent smoothed conditional means with a 0.95 confidence interval. Panels **A**, **B**, and **C** share the same color legend. **D**, Proportion of the different functional categories in each gene class.

### Contrasting distributions of putative virulence factors in the core, dispensable, and private genomes of the three species

Gene-based pangenomes were derived from the graph pangenomes, utilizing the node classification and path of each isolate to categorize the coding regions into four distinct classes: core, dispensable, private genes, and an ambiguous class for genes that did not fit into any of the previous categories. For this classification, genes were labeled as core, dispensable, or private if over 80% of the gene sequence consisted of nodes in that respective class. The ambiguous genes, representing only 5 ± 0.02% in *E. lata*, 3 ± 0.2% in *P. minimum*, and 3 ± 0.1% in *N. parvum*, were not included in further analyses or discussion.

Pangenome modeling helped illustrate the changes in gene numbers as more genomes were incorporated (Figure 3C). *Eutypa lata* and *P. minimum* both exhibited a pronounced slope in the alteration of core, dispensable, and total genes (i.e., the pangenome) with the addition of more genomes. Specifically, in *E. lata*, a stabilization in the change was noticeable after the 10th genome was added. In contrast, for *P. minimum*, this stabilization was not apparent, possibly due to the smaller number of available genomes (Figure 3C). In *N. parvum*, a more moderate and almost linear change in gene numbers was observed, extending up to 16 available genomes. It’s also noteworthy that the number of private genes exhibited minimal change, even in *E. lata* and *N. parvum*, where the models comprised more genomes (Figure 3C). It is interesting to note that *N. parvum* isolates had the largest proportion of genes in the conserved region (93.9 ± 0.1%), compared to 90 ± 0.4 % in *P. minimum* (*P* < 5.78e-06, Wilcox test) and 89.2 ± 0.1% in *E. lata* (*P* < 1.14e-12, Wilcox test). On the other hand, 6.9 ± 0.5% of the genes in *P. minimum* isolates were classified as dispensable or private, 5.5 ± 0.3% in *E. lata*, and, as expected, *N. parvum* had the least proportion of variable genes with 3.3 ± 0.1%. These variable genes had a significantly higher than expected density at the ends of the chromosomes (*P* < 0.1, Chi-squared test), especially in *E. lata* and *P. minimum* (Supplementary figure 3A). Additionally, in *N. parvum*, a significantly larger proportion (Wilcox test, *P* < 0.00005) of variable genes (9.1 ± 0.9% of private and 3.6 ± 0.2% of dispensable) were flanked by TEs when compared to the core genes (0.8 ± 0.03%; Supplementary figure 3B). In *E. lata,* the highest percentage of genes flanked by TEs was observed among the private genes, followed by the core and dispensable genes, while in *P. minimum* there were no significant differences between the gene classes (Supplementary figure 3B).

Notably, *N. parvum* contained the highest proportion of its BGC genes in the core genome, at approximately 95%, followed by *P. minimum* at around 86%, with *E. lata* having the smallest proportion at 82%. These differences reflected an almost double representation of BGCs in the dispensable genomes of *E. lata* (11%) and *P. minimum* (9%), compared to *N. parvum* (4%) (Figure 3D). Overall, genes in BGCs were enriched in the dispensable genome of all the species. Type 1 polyketide synthases (T1PKS) were enriched in *E. lata* (*P* = 2.84e-268, Fisher’s Exact Test), and *P. minimum* (*P* = 3.65e-09, Fisher’s Exact Test), whereas T1PKS/terpene genes were enriched in *N. parvum* (*P* = 6.44e-170, Fisher’s Exact Test). The pattern observed in the BGC genes is almost mirrored in the P450s. However, a divergence is seen when examining the transporters; here, *P. minimum* had the smallest proportion of genes in the core and the largest in the dispensable category. Genes of the Major facilitator superfamily were enriched in the dispensable genome (*P* = 1.17e-06, Fisher’s Exact Test). Another significant finding was the enrichment of secreted CAZymes in the core genome across all three species (Fisher’s Exact Test, *P* = 8.10e-42 in *N parvum*, *P* = 8.10e-42 in *N. parvum*, *P* = 9.55e-17 in *P. minimum*, and *P* = 1.11e-230 in *E. lata*). Only about 1%, 2%, and 4% of secreted CAZymes were in the dispensable genome, with *E. lata* having the lowest proportion, followed by *N. parvum* and *P. minimum*, respectively (Figure 3D). These contrasting distributions within the core and dispensable genomes suggest distinctive adaptive strategies employed by each species.

**Figure 4.**
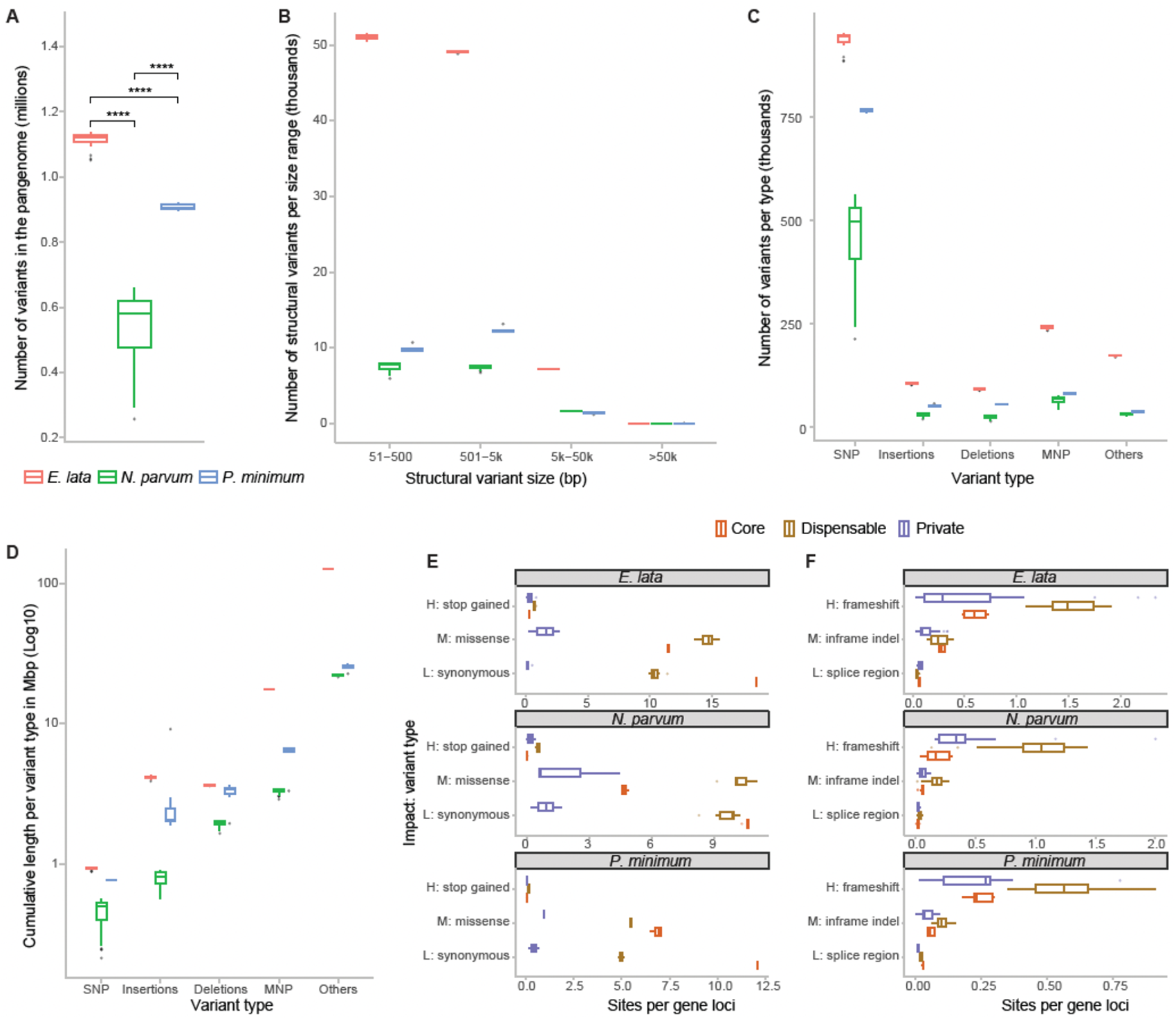
Properties of the genetic variants extracted from the pangenome graph topology. **A**, Total number of variants per species. Significant differences between groups were determined after a Kruskal-Wallis test (*P* < 0.05). **B**, Distribution of variants by size. **C**, Number of variants per type. **D**, Cumulative length of the variants per type. Panels **A**-**D** share the same color legend. **E**, Functional impacts of the SNPs after annotation using SnpEff. The “H” means high, “M” means moderate, and “L” means low. **F**, Functional impacts of the Indels after annotation using SnpEff. The “H” means high, “M” means moderate, and “L” means low. Panels **E** and **F** share the same color legend.

### Analysis of variants in pangenome graphs reveals distinct patterns of SNPs and indels across core and dispensable genes in the three species

The pangenome graphs, built through all-vs-all-pairwise sequence alignments, store information about variants between the genomic sequences used in their construction. We extracted this information using the ‘deconstruct’ command in vg v1.40.0 (43), and subsampled the pangenomes of *E. lata* and *N. parvum* to six genomes, ensuring comparability across the three species. *Eutypa lata* and *P. minimum* exhibited significantly more total variants than *N. parvum*, with respective counts of ∼1.1 million, ∼0.9 million, and ∼0.6 million (Figure 4A). Upon classifying these variants by size and type, the most abundant structural variant group fell within the 50 – 5000 bp range (Figure 4B), with *E. lata* leading, followed by *P. minimum* and *N. parvum*. Though SNPs were the most common variant type (Figure 4C), they accounted for the smallest cumulative length (Figure 4D).

To assess the potential impact of the variants captured within the pangenomes, we annotated all variants using SnpEff 5.1 (44). For this analysis, the total number of genomes per pangenome was utilized, enabling quantitative comparisons within each species, while only qualitative pattern comparisons were made between species. The analysis revealed that low-impact synonymous SNPs (those causing synonymous mutations) were more prevalent in the core genes (Figure 4E). In general, most SNPs in the core genes have low impact as expected in genes that are conserved across isolates. Moreover, in *E. lata* and *P. minimum*, low-impact synonymous SNPs occurred roughly twice as often in core genes compared to dispensable genes. In *N. parvum*, however, the frequency was nearly the same in both gene categories. Conversely, high-impact SNPs (those introducing premature termination codons) and moderate-impact SNPs (those causing missense mutations) were generally more common in dispensable genes. Concerning indels (Figure 4F), high-impact frameshift mutations were more commonly found in dispensable genes. Moderate-impact in-frame indels also tended to be more frequent in dispensable genes, although in *E. lata*, their occurrence was nearly equal in core genes. Low-impact indels, which cause changes in the splice region, exhibit a similar frequency across all gene classes

### Biosynthetic gene cluster genes in the dispensable genomes of *E. lata* and *P. minimum* are under positive selection

Gene selection patterns in the isolates of three species were analyzed by calculating the ratio of non-synonymous (dN) to synonymous (dS) mutations using the YN00 model in the PAML package v.4.9 (47). The analysis revealed that the majority of genes (∼81 – 86%) are under negative or purifying selection (dN/dS < 1). Positive selection (dN/dS > 1) was identified in only 1.8% to 2.2% of the genes (Figure 5A). Notably, genes under positive selection were enriched in the dispensable genome of the three species (Fisher’s Exact Test*, P* = 3.28e-06 in *E. lata*; *P* = 0.00056 in *P. minimum*; *P* = 0.0023 in *N. parvum*). Moreover, the proportion under positive selection was almost double in the dispensable than in the core genes, representing approximately 5% of the dispensable genes in *E. lata* and approximately 4% in *N. parvum* and *P. minimum* (Figure 5B).

Additionally, the analysis revealed specific patterns in the selection of certain gene categories. No gene annotated as a CAZyme was under positive selection in *E. lata*, whereas in *N. parvum* and *P. minimum*, approximately 0.6% and 1% of the annotated CAZymes were under positive selection, respectively. For BGC genes, approximately 2.2% and 2.7% were under positive selection in *E. lata* and P*. minimum*, respectively, compared to only 1% in *N. parvum* (Supplementary table 3). Of the BGCs genes under positive selection, two and four are dispensable genes in *P. minimum* and *E. lata,* respectively, and 12 are part of the core genes in the two species. Among the genes annotated as transporters, 0.6% were under positive selection in both *E. lata* and *N. parvum*, compared to 1% in *P. minimum*. Of these transporters, 28, 17 and nine genes were in the core genomes of *P. minimum*, *N. parvum* and *E. lata,* respectively, compared to three, one and four in the dispensable genomes. Lastly, of the P450 genes, 0.3% in *N. parvum* were under positive selection, compared to 0.6% in *E. lata* and 1.4% in *P. minimum*. Most P450 genes under positive selection were in the core genome. These findings highlight the differential selection pressures on various functional gene categories across the three species.

**Figure 5.**
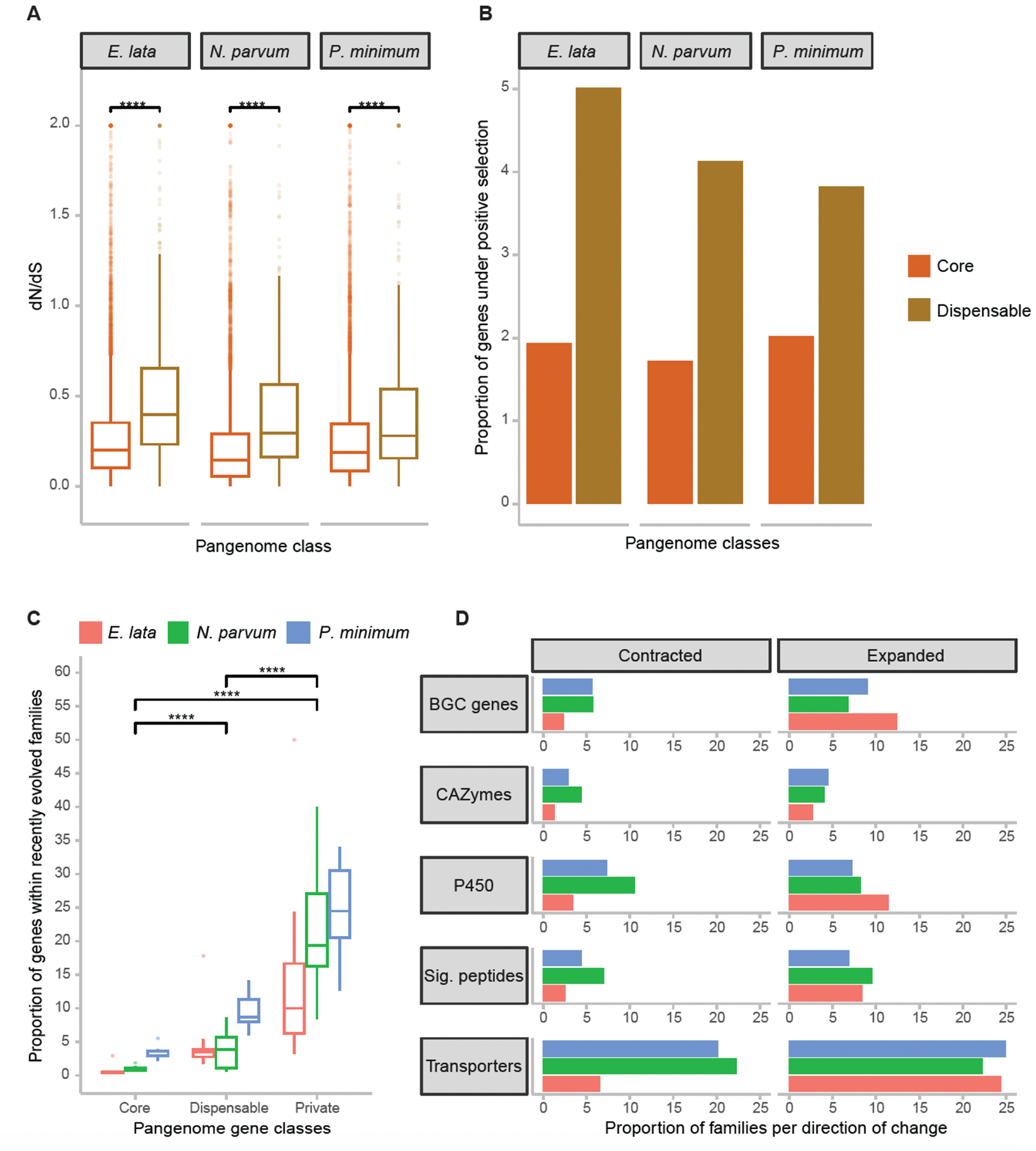
Gene evolution patterns in the pangenomes of the species in study. **A**, dN/dS ratios of the genes in the different pangenome classes. Significant differences were determined using a two-tailed Student’s t-test (*P* < 0.05). **B**, Proportion of genes in each pangenome class under positive selection. **C**, Expanded/Contracted gene families per pangenome class. Significant differences were determined after a Kruskal-Wallis test (*P* < 0.05). **D,** Proportion of expanded and contracted families annotated with the different putative virulence factors. Panels **A** and **B** share the same color legend. Panels **C** and **D** share the same color legend.

### Analysis of rapidly changing gene families and functional categories reveals distinct evolutionary patterns in the three species

An analysis of gene family evolution, conducted using the CAFE5 pipeline (48), identified expanded and contracted gene families among isolates of each species: 650 in *P. minimum*, 549 in *E. lata*, and 472 in *N. parvum* (Supplementary figure 4-6). Notably, the genes of the private genomic regions were most affected by these changes (Figure 5C), with 24 ± 3% of the private genes in *P. minimum*, 22 ± 4% in *N. parvum*, and 13 ± 2% in *E. lata* being part of these dynamic families. Dispensable genes were the next most affected, with proportions ranging from ∼9 to 4%, while core genes were least affected, with values ranging from ∼3 to 0.5%.

Among the recently evolved families in *E. lata*, approximately 13% displayed BGC genes or P450 genes expanding, whereas only ∼2% were contracting (Figure 5D). The BGC genes in expansion represented 21.5% of the dispensable genes in *E. lata* genomes, and only 2.6% of the total core genes. On the other hand, P450 contracting families in *N. parvum* represented 11% of the recently evolved families in the species. Concerning CAZymes, *E. lata* exhibited the least changing families, followed by *P. minimum*, while the most significant change was observed in *N. parvum*, with a combined 9% of the total families undergoing expansion and contraction containing CAZyme genes. The expanding CAZymes represented 4.2% and 2.8% of the total core and dispensable genes, respectively, in *N. parvum* genomes. Interestingly, *N. parvum* had the highest proportion of families annotated with signal peptides undergoing change (∼17%), compared to *E. lata* and *P. minimum* (∼11%).

Transporters emerge as the most prominent functional category within these evolving gene families. In each species, over 20% of the total families contain transporter genes undergoing expansion. A similar percentage is noted for contraction in *N. parvum* and *P. minimum*, but in *E. lata*, only ∼7% of the total families are contracting (Figure 5D). The contracting transporter genes represent 19.0% and 14.6% of the core genes of *P. minimum* and *N. parvum* genomes, respectively, compared to only 6.7% in *E. lata* genomes.

To analyze specific functions within these broader functional categories, we conducted an enrichment test of the genes in expanded and contracted families. The test revealed that in *E. lata*, genes of the T1PKS cluster were highly enriched in the expanded families (*P* = 2.24e-17, Fisher’s Exact Test; Supplementary figure 7). Additionally, CYP149, known for its polyketide-synthase functions, was also enriched in the expanded family (*P* = 5.94e-21, Fisher’s Exact Test). Conversely, *P. minimum* exhibited CYP53 with benzoate 4-monooxygenase activity enriched in the contracted families (*P* = 1.67e-40, Fisher’s Exact Test; Supplementary figure 8). The expanded and contracted families of this species also showed enrichment in the transporters 2.A.1 (Major Facilitator Superfamily, *P* = 1.87e-58, Fisher’s Exact Test), which, among other roles, have been involved in resistance to oxidative stress and fungicides in some pathogenic fungi (49, 50). Lastly, the expanded and contracted families in *N. parvum* are enriched in glycosyltransferases (GT1_153) (*P* = 1.08e-16, Fisher’s Exact Test, Supplementary figure 9), which have been associated with virulence in some pathogenic fungi (51, 52).

## Discussion

Virulence variations among grapevine trunk pathogen species are frequently observed, often attributed to the complex and adaptable nature of fungal genomes (53–55). This complexity can confound traditional comparative genomics methods, as relying on a single representative for species characterization may lead to biased conclusions, especially if the chosen isolate lies at an extreme of the virulence spectrum. Recently, reference-free sequence-based pangenomes have emerged to address this challenge (46, 56). By encompassing a collection of isolates across the virulence spectrum, pangenomes offer a more comprehensive view of a species’ genomic and functional characteristics. This approach enhances our understanding of virulence variations, providing a more objective depiction of the species’ functions (57–59).

In this study, we assembled and annotated the genomes of fifty isolates of the three critical pathogens associated with Eutypa dieback, Botryosphaeria dieback and Esca. The average assembly size, completeness and gene predictions of the species are consistent with previously published genomes (3, 5, 6, 20, 60). Prior to the construction of the three pangenomes, we validated the completeness of the fifty assemblies by structural and gene-content comparisons with chromosome-scale assemblies based on long-read sequencing, including a new one for *E. lata* assembled as part of this study. We then used whole pair-wise genome alignments to produce the first fungal reference-free nucleotide-scale pangenome graphs. This method avoids the bias inherent in reference-based approaches and, in contrast to the more common gene-based pangenomes, incorporates non-coding regions, regulatory elements, and intergenic regions. This inclusion allows for a more comprehensive understanding of the genomic architecture and functional elements (61).

In the three pangenomes, the private fractions of the genomes contain the largest proportion of repeats, followed by the dispensable and core regions. Transposable elements like the Gypsy encountered in our analysis, contribute to genetic variation by inducing mutations, rearranging genomic structures, and influencing gene expression (62–64). In the private fractions of the genomes, which often contain genes that confer specific adaptations or characteristics, the presence of repeats may facilitate rapid evolutionary changes, leading to differentiation between individual genomes within a species (65).

Comparisons of the three pangenomes also revealed important differences between the three species. The first notable difference observed was that *E. lata* possesses the largest number of nodes in its pangenome graph. This could be indicative of a more complex genomic structure, encompassing various genomic variants, as suggested by previous research (17, 20, 66). While the utilization of more isolates in building *E. lata*’s pangenome might contribute to this observation (67), the similarity in node complexity between *P. minimum* and *N. parvum*—each with approximately four million nodes—supports that the number of genomes used is not the sole factor for greater node complexity. Furthermore, over half of *E. lata*’s total pangenome length consisted of variable regions (either dispensable or private). This contrasts with the lower percentages observed in *P. minimum* and *N. parvum*. This finding further reinforces the idea that *E. lata* exhibits a higher complexity of genomic variants, distinguishing it from the other two species.

The analysis of the gene-based pangenomes showed a predictable pattern where the total and dispensable genes increase with the addition of more genomes, while the core genes decrease. However, the rate of these changes varies significantly among the three species, a phenomenon that is particularly pronounced in the gene-based model, but also discernible in the sequence-based model. One possible explanation for *E. lata* and *P. minimum* exhibiting steeper changes is that these two species produce sexual spores (5, 20, 66, 68). As a result, the genomic recombination process likely leads to a greater number of genomic variants between isolates. Conversely, *N. parvum*, which spreads in the form of asexual spores (69, 70), is expected to generate fewer genomic variations. This could account for the less pronounced alterations in the pangenome classes, as additional genomes are incorporated. Moreover, the higher density of variable genes in the terminal and subterminal regions of the chromosomes may help explain the differences between species. These regions, known for their greater genomic instability (25, 71), also serve as recombination hotspots in fungi (72). Additionally, the bigger proportion of dispensable and private genes (compared to core genes) flanked by TEs in *N. parvum*, suggest that these elements could be one of the factors introducing the gene variation in a species with limited sexual recombination (65). It is also important to highlight that the number of isolates available for *P. minimum* pangenome may be insufficient to accurately predict the trend of the pangenome classes. Nevertheless, it is evident that the behavior of the classes in *P. minimum* is closer to what we observe in the first six genomes of *E. lata* than those in *N. parvum*.

The larger number of genomic variants in the pangenome graphs for *E. lata* and *P. minimum* supports the notion that sexual reproduction is one of the main drivers of intraspecific diversity. When evaluating the number of structural variants by size, these two species exhibit larger numbers across all sizes. However, *E. lata* shows strikingly higher numbers than *P. minimum*, particularly in variants ranging from 50 bp to 50 kbp. This difference may be attributed to the higher proportion of repetitive content in *E. lata* genomes compared to *P. minimum*, contributing to greater genomic variation (64, 65).

Not all the variants are expected to have same impact on the isolate phenotypes. Therefore, predicting the type of change generated by these variants and their impact is crucial to better describe the genomic variability of the species (73, 74). Our results are consistent with previous studies (75) indicating that low-impact variants are generally more abundant in the core genome, whereas high and medium-impact variants tend to be more prevalent in the dispensable regions of the genome.

The analysis of the ratio of non-synonymous to synonymous mutations revealed interesting evolutionary patterns. Most genes exhibited dN/dS values below 0.9, indicating minimal change in protein sequence, a result consistent with the literature (76, 77). In the context of the pangenome, our analysis revealed that the proportion of genes exhibiting signatures of positive selection was over twice as prevalent in variable regions as in conserved ones. Notably, many of the genes under positive selection in the dispensable regions of the pangenomes were BGC genes in both *E. lata* and *P. minimum*, corroborating previous findings of intraspecific variation affecting BGC genes (5, 20).

Slight variations in the genes of biosynthetic gene clusters (BGCs) can lead to multiple variants of secondary metabolite backbones, potentially altering their functions (78–80). *E. lata* and *P. minimum* are great examples of this phenomenon. In *E. lata*, compounds like eutypine, eutypinol, O-methyleutypine and many others, share a common backbone but exhibit varying toxicities within and between hosts, with eutypine as highly toxic, compared to its non-toxic derivative eutypinol (10, 81–83). Furthermore, apart from the extensive array of secondary metabolites in *P. minimum*, diverse virulence modes of action have been documented. These range from exopolysaccharides and polypeptides, causing varying degrees of disruption to water transport (10, 84, 85), to quinones and melanins, which shield the fungi against reactive oxygen species produced by plants as part of their defense mechanisms (10, 86–88). This suggests that changes in BGC gene content, and the corresponding secondary metabolites they produce, may be instrumental in adapting to new environmental conditions and could underlie a significant portion of intraspecific variations in virulence levels.

Additionally, among the three species, *N. parvum* showed the highest number of CAZymes under positive selection, whereas *E. lata* had no CAZymes with ratios indicative of positive selection. While it is essential for the majority of grapevine trunk pathogens to effectively breach the plant cell wall in order to successfully establish colonization, it has been shown that *N. parvum* is known for being an efficient wood degrader that relies heavily on the diversity and plasticity of its enzymatic arsenal to efficiently degrade non-structural and structural complex polymers of the plant host cell walls and access cellular content for nutrition (89). The recent expansion of CAZymes (3), and their significantly augmented expression in the presence of wood (6), suggest that this group of enzymes contributes significantly to the virulence variability observed in *N. parvum*.

We also observed differences in transporters, P450, and peroxidases, indicating that although all putative virulence factors are vital for the three species, they depend on modifications in different virulence factors to adapt to their specific conditions (3). It is worth noting, as emphasized by other researchers (90–92), that in microbial populations, genes under positive selection may have dN/dS values less than one. Therefore, the results presented here might be an underestimation of the true number of genes under positive selection.

By leveraging the genetic information captured in the pangenomes, we have identified distinct patterns of expansion and contraction of gene families across these species. A greater proportion of genes in the variable regions of the pangenome exhibited rapid changes, with the core regions being the least influenced by the expansion and contraction of these gene families. The evolutionary trends revealed in our findings align with that of prior research (75), in which BGCs gene families are undergoing frequent changes in *E. lata* and *P. minimum*. Moreover, our results corroborated earlier observations indicating dynamic changes in the gene families associated with cellular transporters in *P. minimum* (3, 5). On the other hand, *N. parvum* stands out as the species displaying the highest count of gene families annotated with CAZyme functions undergoing rapid evolution, contradicting previous assertions where *E. lata* was attributed with more significant numbers than *N. parvum* (3). This inconsistency could potentially arise from the fact that the previous study reached its conclusions based on a single isolate per species. The enrichment of specific gene functions within these rapidly changing families is noteworthy. These genes not only help elucidate the evolutionary patterns among species, but also highlight the differences within each species.

By incorporating a comprehensive view of the genomic variation, which a single reference genome of a species cannot achieve, the pangenomes developed in this study are valuable tools for future forward genetic analyses like pangenome wide-association analysis (panGWAS; (75)) approaches, aimed at uncovering the genetic basis of specific pathogenic and virulence-related phenotypes, or metatranscriptomic profiling of gene expression of fungal populations during natural infections(93).

## Methods

### Sampling and genome assembly

All fungal isolates utilized in this study were obtained from wood of symptomatic hosts following the protocol of previously described by Baumgartner *et al*. (94).

*Eutypa lata* isolate Napa209 (or EN209) was grown in Potato Dextrose Broth at 150 rpm for 7 days. In preparation for DNA extraction, the mycelium was filtered, washed with sterile water, and frozen in liquid nitrogen. The frozen mycelium was ground with a TissueLyser II (Qiagen) using a 50-ml stainless-steel jar (Retch). High molecular DNA extraction was based on the method of Stoffel et al. (95), using 4 g of ground mycelium. The molecular weight was evaluated with the Pippin pulse electrophoresis. DNA was fragmented using the Megaruptor (Diagenode) and again evaluated with pulse electrophoresis. A total of 15 μg of fragmented, high molecular weight DNA was then used as input for library preparation, with the SMRTbell Template Prep Kit 1.0 (Pacific Biosciences) following manufacturer instructions. Ligated products were size-selected for fragments larger than 12 kb, using Blue Pippin (Sage Science). Finished libraries were submitted for sequencing in the PacBio RS II platform, using the Pacbio P6-C4 chemistry (DNA Technologies Core Facility, University of California, Davis, California, USA).

The total genomic DNA of all 50 isolates was also extracted following the CTAB protocol as presented in the methods of Morales-Cruz et al. (3). Illumina DNA libraries were prepared following the protocol in Morales-Cruz et al. (3). After adapter ligation, libraries were size selected to 400–600 bp using a double-sided size selection with Ampure XP magnetic beads (Beckman Coulter, United States). Sequencing was carried out on an Illumina HiSeq4000 machine at the DNA Technologies Core at UC Davis. Paired-end reads of 150 bp or 100 bp in length were generated for each isolate.

SMRT reads were assembled using HGAP v.3.0 (96). Reads were filtered based on the following parameters: minimum sub-read length = 1,000 bp; minimum polymerase read quality = 0.8; and minimum polymerase read length = 500 bp. The minimum seed read length for assembly was set at 6,000. Assembly polishing was carried out with Quiver, using only unambiguously mapped reads. To estimate the error rate, Illumina paired-end reads of the isolate were mapped using Bowtie2 v.2.2.6 (97), PCR and optical duplicates were removed with Picard tools (v.2.0.1; http://broadinstitute.github.io/picard/).

Illumina raw reads were quality trimmed with Trimmomatic v0.36 (98) following methods previously described (7). SPAdes v3.13.0 (99) was used to assembled the genomes with the careful option and automatic read coverage cutoff after optimizing the multiple kmer combination.

The repeats library 20160829 was used with RepeatMasker open-4.0.6 (100) to annotate and mask the repeats in all genomes. The gene model prediction was performed with RNAseq data mapped to the genomes using the BRAKER1 pipeline (101) which combines GeneMark-ET (102) and Augustus (103)

Assembly completeness was assessed using the Benchmarking Universal Single-Copy Orthologs analysis (BUSCO v. 5.4.2; (104). Additionally completeness was evaluated with GMAP v2019-09-12 (105) using the default parameter and counting the proportion of the genes in the PacBio assembled genomes with minimum 90% covered by the Illumina assembled genomes. Additionally the percentage of genes in colinear blocks was determined by MCScanX (106) using default parameters.

### Functional annotation

The annotation of conserved protein domains was done with the Pfam database (107). The annotation of putative virulence factors was assigned based on the databases and parameters presented in Supplementary table 4. CAZymes were annotated with the dbCAN3 (108). The signal peptides were predicted using SignalP 5.0 (45). The proteins with annotation in both databases were annotated as secreted CAZymes. Secondary metabolite clusters were annotated using antiSMASH 6.0.1 (36). Peroxidases were annotated using a specialized database for fungi called fPoxDB (109). CYPED v6.0 was used to annotate the Cytochrome P450 proteins (39). Lastly, the proteins involved in transportation functions were annotated using the TCDB (43).

### Nucleotide level reference-free pangenome assembly

The seq-based pangenome graph was built using the tools in the PGGB pipeline(46). The assembled genomes were indexed by samtools v1.7 (110, 111) and aligned in a pairwise manner with wfmash (https://github.com/waveygang/wfmash) using the optimized parameters “-p 90 -s 4000 -n 1”. The alignments were concatenated per species and processed with seqwish v0.7.3 (112) to produce the pangenome graphs with the optimized kmer size of 47. Finally, to avoid node ambiguities and keep the nodes sorted, smoothxg (https://github.com/pangenome/smoothxg) was used in two rounds. The first round used the following common parameters: “-K -X 100 -I 0.85 -R 0 -j 0 -e 0 -l 4001 -P “1,4,6,2,26,1” -O 0.03 -Y 1700 -d 0 -D 0 -V” with the corresponding -w for each species. The parameters for the second were the same changing the -l parameter to 4507 and therefore the -w parameter for each species. These options were selected based on the default target poa lengths (-l) of 4001 and 4507 in the PGGB command. For each species, the nodes and sequences were classified into the core, dispensable, or private. Nodes were classified as core if they were shared by all the genomes in a pangenome, dispensable if the nodes were shared by more than one genome but not all, and private nodes when they were exclusively present in a single genome.

For the gene-based pangenome inference, the node information was intersected with the gene annotation file using BEDtools v2.29.1 (113). The genes were classified as core, dispensable or private when it sequences were covered more than 80% by nodes in that particular class. The genes not falling in any group were classified as ambiguous.

### Annotation of genes in terminal chromosomal regions and genes flanked by TEs

All complete chromosomes of long reads assemblies were divided in 10% windows. The location of the genes in each windows were calculated with BEDtools intersect v2.29.1 and classified in the different pangenome classes. Chi-squared test was used to determine if the observed frequency distribution fits the expected distribution.

To identify the genes flanked by TEs we selected 500bp at each side of the genes of all the isolates and intersected them with the TEs annotation. Genes with at least one TE in any of the flanking regions were counted. The proportion of the genes affected by TE were calculated per pangenome class and a Pairwise Wilcox test was used to determine significant differences.

### Genomic variants extraction and annotation

Genomic variants were extracted and categorized from the pangenome graph using vg deconstruct v1.40.0 (114) and bcftools v1.8 (111) following the protocol in https://github.com/noecochetel/North_American_Vitis_Pangenome (75). The functional impact of the different types of variants was predicted SnpEff v.5.1 (74).

### Phylogenetic relationship within species

The SNPs obtained from the pangenome graph were filtered to remove variants in repetitive regions. Since the haplotype of each isolate was recorded for each SNP position a synthetic sequence alignment was produced by concatenation of these positions. These alignments were parsed with Gblocks v0.91b (115), with default parameters per species. Next, the phylogenetic analysis was performed using RAxML-NG v0.9.0 (116) with the option --opt-model to optimize the evolutionary model in function of the data. We applied the Maximum Likelihood (ML) method with the optimized evolutionary model GTR, using ten random starting trees, a bootstrapping of 100 replicates and the Gblocks parsed alignments.

### Patterns of selection, expansion and contraction of gene families

To evaluate the dN/dS ratios of all genes, the SNPs were obtained by mapping the trimmed reads of all the isolates to CDS of the PacBio assembled genomes of each species using bwa-0.7.12 (117). Picard tools v.2.0.1 (http://broadinstitute.github.io/picard/) was used to remove duplicates. The resulting bam was used as input for GATK v4.0.12.0 HaplotypeCaller (118) to call the variants. Only SNPs were kept using bcftools v1.8. These SNPs were incorporated into the reference CDS with GATK FastaAlternateReferenceMaker, generating orthologues for each gene of the reference in all isolates. ParaAT 2 (119) was used to generate the protein and CDS alignments of the orthologues genes. The dN/dS ratios were calculated with the YN00 model in the PAML package v4.9 (47). For this analysis, ratios below 0.9 were considered under negative selection, above 1.1 were considered under positive selection and in between as neutral. The Fisher’s exact test was used for the enrichment analysis of these genes.

To study the patterns of gene family evolution, the predicted proteins of the isolates of each species were compared to each other using OrthoFinder v.2.5.4 (120). For this analysis we included related species as outgroups. For *P. minimum* and *E. lata* we used *Fusarium oxysporum* Fo47 v2.0 (121) and *F. solani* FSSC 5 v1.0 (122). For *N. parvum* we included *Alternaria alternata* MPI-PUGE-AT-0064 v1.0 (122) and *A. brassicicola* (123). The sequences of each single-copy gene orthogroup were aligned using MUSCLE v.5.1 (124). Alignments were concatenated and parsed with Gblocks v.0.91b with default parameters. The LG evolutionary model was selected using ModelTest-NG v.0.1.7 (125).

The clock-calibrated tree was crated with BEAUti and BEAST v.1.10.4 (126). The starting tree was constrained to the SNP tree topology with monophyletic partitions. Calibration points were set for different groups: The crown age of the Sordariomycetes group was set to ∼233 mya (127, 128). The Nectriaceae group age was set to ∼97 mya (128). For the Dothidiomycetes group the age was set to ∼366 mya (123, 127, 129). The divergence age of the Alternaria species was set to ∼10 mya (123). Then, six independent Markov chain Monte Carlo (MCMC) chains, each consisting of 1,000,000 generations were run. The LG substitution model, with four Gamma categories, a strict clock, and the Birth-Death Model was used. Sampling was performed every 1,000 generations. The resulting log and tree files were combined using LogCombiner v.1.10.4 (126), and the maximum clade credibility tree was generated using TreeAnnotator v.1.10.4 (126) with a burn-in of 10,000 generations.

The final estimation of family expansion and contraction was done with diamond v0.9.24.125 (130), mcl v14-137 (131) and CAFE v.5.0 (48) as described in (75). The estimated lambda parameter for *P. minimum* was 0.00416, for *E. lata* was 0.00251, and for *N. parvum* was 0.000975.

### Data visualization and statistical analysis

RStudio v.2023.03.0.386 (132) running R v.4.2.2 (133) was used to analyze the data and to produce all the figures mostly using the package tidyverse v.2.0.0 (134). Phylogenetic trees were drawn using the R package ggtree v.3.4.4 (135) and Figtree v1.4.4 (136).

## Data availability

The sequencing data generated for this project is available at NCBI (BioProject PRJNA1009151). The sequencing data of the previously published chromosome scale assemblies of *P. minimum* Pm1119 (5) and *N. parvum* UCD646So (6) can be retrieved from NCBI under BioProject PRJNA421316 and PRJNA321421, respectively. The NCBI BioProject accession for sequencing data previously published and used in this study can be seen in Supplementary table 1. All genome assemblies and gene models are publicly available at Zenodo (doi: 10.5281/zenodo.8310403).

## Acknowledgments

This work was supported by the USDA, National Institute of Food and Agriculture, Specialty Crop Research Initiative (grant 2012-51181-19954) and by the American Vineyard Foundation (grant 1798). DC was also partially supported by the Louis P. Martini Endowment in Viticulture and the UC Davis Chancellor’s Fellow Award.

